# Myosin-II mediated traction forces evoke localized Piezo1 Ca^2+^ flickers

**DOI:** 10.1101/294611

**Authors:** Kyle L. Ellefsen, Jesse R. Holt, Alice Chang, Jamison L. Nourse, Janahan Arulmoli, Armen Mekhdjian, Hamid Abuwarda, Francesco Tombola, Lisa A. Flanagan, Alexander R. Dunn, Ian Parker, Medha M. Pathak

## Abstract

Piezo channels transduce mechanical stimuli into electrical and chemical signals, and in doing so, powerfully influence development, tissue homeostasis, and regeneration. While much is known about how Piezo1 responds to external forces, its response to internal, cell-generated forces remains poorly understood. Here, using measurements of endogenous Piezo1 activity and traction forces in native cellular conditions, we show that actomyosin-based cellular traction forces generate spatially-restricted Ca^2+^ flickers in the absence of externally-applied mechanical forces. Although Piezo1 channels diffuse readily in the plasma membrane and are widely distributed across the cell, their flicker activity is enriched in regions proximal to force-producing adhesions. The mechanical force that activates Piezo1 arises from Myosin II phosphorylation by Myosin Light Chain Kinase. We propose that Piezo1 Ca^2+^ flickers allow spatial segregation of mechanotransduction events, and that diffusion allows channel molecules to efficiently respond to transient, local mechanical stimuli.

## INTRODUCTION

Cells both detect and generate mechanical forces, and integrate mechanical information with genetic and chemical cues to shape organismal morphology, growth, and homeostasis. Mechanical forces are transduced into biochemical signals by specialized proteins. Among these, mechanically-activated ion channels provide unique features: sub-millisecond response to mechanical stimuli, high sensitivity, large dynamic range, spatial coding of mechanical stimuli, and the ability to temporally filter repetitive stimuli ^1^.

Piezo channels were recently identified as a new family of excitatory mechanically-activated channels ^2,3^. Due to their permeability to Ca^2+^ and other cations, Piezo channel activity generates chemical as well as electrical signals in response to mechanical stimuli, allowing them to regulate a wide variety of cellular processes. Indeed, Piezo1 has emerged as an important player in vascular development ^4,5^, stem cell differentiation ^6,7^, epithelial homeostasis ^8^, bladder mechanosensation ^9^, erythrocyte volume regulation ^10^, cell migration ^11–13^, vascular smooth muscle remodeling ^14^, cartilage mechanics ^15,16^, blood pressure regulation ^17^ and exercise physiology ^18^. The global knockout of Piezo1 is embryonic lethal ^5^, and mutations in the channel have been linked to diseases such as dehydrated hereditary stomatocytosis ^19–22^, colorectal adenomatous polyposis ^23^, generalized lymphatic dysplasia ^24,25^ and malarial parasite infection ^26^. Thus, understanding how Piezo1 functions is critical for deciphering its diverse physiological roles.

Studies on Piezo1 have largely focused on transduction of “outside-in” mechanical forces, i.e. forces arising from stimuli such as poking, negative suction pulses, shear flow, osmotic stress and displacement of the substrate ^2,4,27– 30^. However, cells also actively generate mechanical forces ^31^; for example, cells use Myosin II-generated traction forces for probing the stiffness of the extracellular matrix. Cell-generated traction forces serve as key regulators of cell signaling and function, modulating cell migration, wound healing, cancer metastasis, immune function, and cell fate ^32^. For example, we previously reported that Piezo1 activation is a key mediator of mechanosensitive lineage choice in human neural stem/progenitor cells (hNSPCs), and that activation of Piezo1 in this system required cell-generated traction forces ^7^. Despite its clear physiological importance, the mechanisms by which cell-generated mechanical forces act to activate Piezo1 remain essentially unknown, either in hNSPCs or any other cell type. Uncovering how traction forces activate Piezo1 is vital to understanding the channel’s role in stem cell fate ^7^, cell migration ^13,33^, and cancer ^12,34^.

Piezo1 activation in cells is typically measured by patch clamp assays that drastically affect the native environment of Piezo1, disrupt cellular composition and cytoskeletal dynamics, and provide no spatial information as to where channels are activated. An alternative, non-perturbing method to monitor activation of Piezo1 channels is imaging Ca^2+^ flux through the channel ^7,35^. Using this approach in hNSPCs, we found that traction forces elicit discrete, local, and transient Ca^2+^ microdomains or “flickers” from endogenous Piezo1 channels in the absence of externally-applied mechanical forces ^7^.

Here we examine the spatial regulation of Piezo1 by traction forces by imaging Piezo1 Ca^2+^ flickers, traction forces and the Piezo1 protein localization in live cells. Imaging Piezo1 Ca^2+^ flickers at submicron-scale spatial and millisecond-scale temporal resolution while manipulating or measuring traction forces reveals several key findings. Although Piezo1 channels are motile in the plasma membrane and are widely distributed across the cell, flicker activity is only enriched in the vicinity of force-producing adhesions. Moreover, Piezo1 Ca^2+^ flickers are triggered by activation of Myosin II through phosphorylation by Myosin Light Chain Kinase (MLCK) but not by Rho-associated protein kinase (ROCK). In light of recent evidence demonstrating that membrane tension gates Piezo1 ^27,28,36^, our studies suggest that cellular traction forces generate local increases in membrane tension that activate Piezo1 within spatial microdomains. The spatial specificity of Piezo1 Ca^2+^ flickers elicited by traction forces may serve to localize downstream biochemical signaling, allowing spatial segregation of mechanotransduction events. We further propose that Piezo1 channel mobility allows a small number of channels to explore large areas of the cell surface, and hence respond to both unpredictable external forces as well as to hotspots of cell-generated traction forces.

## RESULTS

### Piezo1 generates Ca^2+^ flickers

We previously reported Ca^2+^ flickers observed by Total Internal Reflection Fluorescence Microscopy (TIRFM) imaging of hNSPCs in the absence of external mechanical stimulation ^7^. These Ca^2+^ flickers were substantially reduced following siRNA-mediated Piezo1 knockdown, indicating that they were largely produced by Piezo1 activity ^7^. We extended the finding to Human Foreskin Fibroblasts (HFFs) and to Mouse Embryonic Fibroblasts (MEFs). Like hNSPCs, both cell types showed Ca^2+^ events in the absence of external mechanical stimulation (Fig. 1). A CRISPR knockout of the Piezo1 gene in HFFs showed 82% reduction in Ca^2+^ flickers compared to wildtype cells (Fig. 1B, Movie M1). Residual Ca^2+^ events in Piezo1 KO HFFs persisted in the absence of extracellular Ca^2+^ (Fig. S1), suggesting that these are produced by liberation of Ca^2+^ from intracellular stores, rather than from other plasma membrane channels. MEFs derived from constitutive Piezo1 knockout mice ^5^ showed 94% lower occurrence of Ca^2+^ flickers compared to MEFs from wildtype littermate embryos (Fig. 1C, Movie M2). Taken together, we provide evidence that a large majority of Ca^2+^ flickers at the cell-substrate interface in hNSPCs ^7^, HFFs, and MEFs derive from Piezo1 activity, and therefore refer to them as “Piezo1-dependentCa^2+^ flickers” or “Piezo1 Ca^2+^ flickers”.

**Figure 1.**
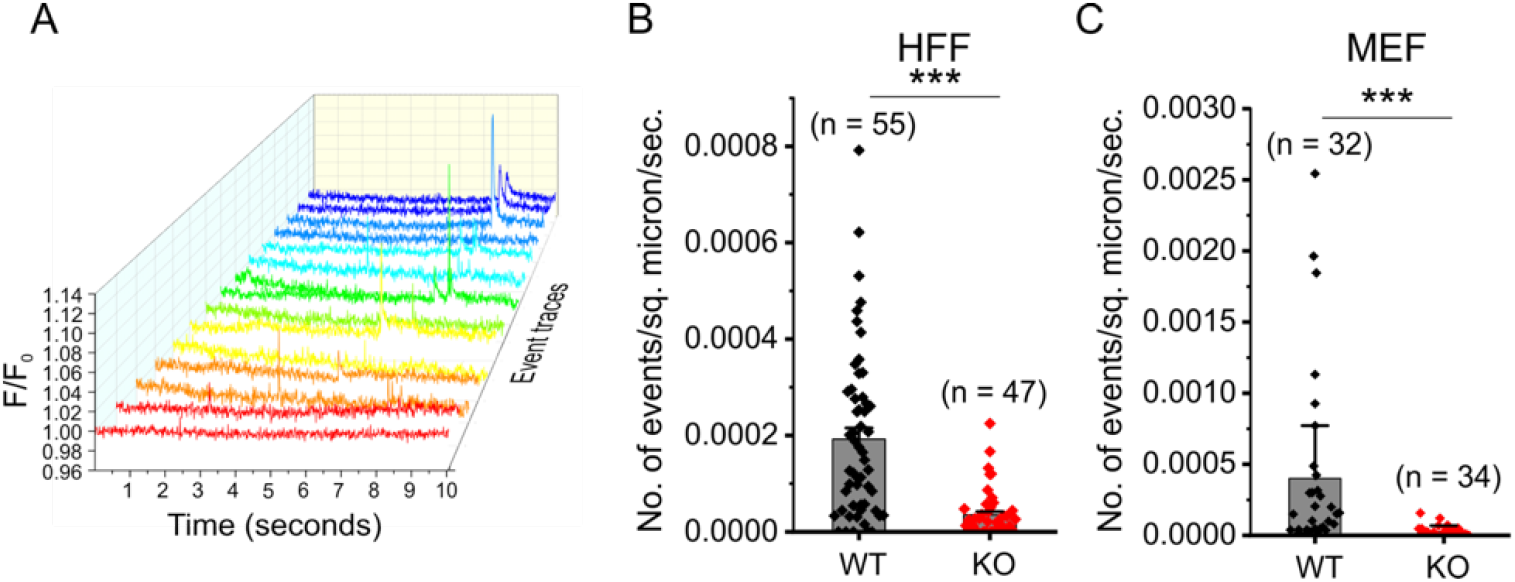
Piezo1 channels generate Ca^2+^ flickers. **A.** Representative Ca^2+^ flickers recorded from HFF cells. Traces show fluorescence ratio changes (ΔF/F_0_) from multiple regions of interest of a video, plotted over time and in different colors for clarity. **B.** Frequency of Ca^2+^ flickers in Wildtype HFFs and in Piezo1knockout HFFs. Bars denote mean ± SEM and each point represents flicker frequency in an individual video. **C.** Frequency of Ca^2+^ flickers in Piezo1-knockout MEFs and in Wildtype MEFs derived from littermate embryos. n values for panels B and C denote number of videos (i.e. unique fields of view, each composed of one or more cells) from three different experiments. *** denotes p < 0.001 by Kolmogorov-Smirnov test. See also Fig. S1 and Movies M1 and M2.

### Super-resolution localization of Ca^2+^ flickers

To examine the spatial relationship of Piezo1-dependent Ca^2+^ flickers relative to hotspots of traction forces, we developed a technique for automated localization of Piezo1 Ca^2+^ flickers at super-resolution levels (Fig. 2). This approach is an improved version of our algorithm for automated detection and quantitation of local Ca^2+^ signals ^37^, implemented as a plugin for the general purpose image processing software Flika (http://flika-org.github.io). The algorithm uses a clustering method ^38^ to group suprathreshold pixels into Ca^2+^ events, improving the unbiased detection and segregation of signals (see Methods for further details).

**Figure 2.**
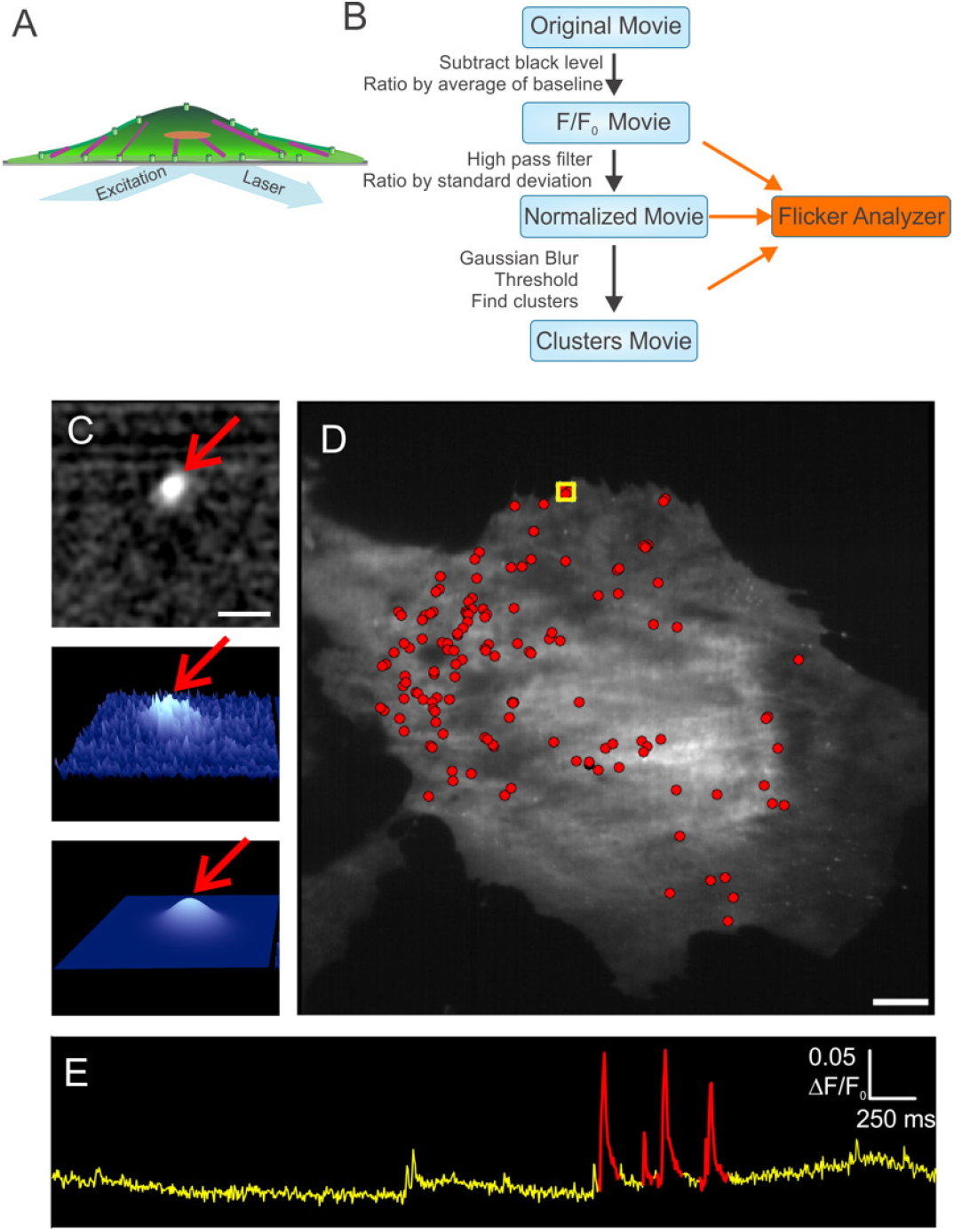
Automated detection and super-resolution localization of Piezo1-dependent Ca^2+^ flickers. **A.** Piezo1 Ca^2+^ flickers are acquired by Ca^2+^ imaging with Total Internal Reflection Fluorescence Microscopy (TIRFM). **B.** Flowchart of the algorithm. The original movie is processed to subtract camera black level and then divided by the average of the first ~100 frames to produce a F/F_0_ ratio movie. The ratioed movie is spatially and temporally filtered to increase the signal-to-noise ratio. A clustering algorithm groups supra-threshold pixels into flicker events. **C.** For every event detected, a 2D Gaussian fit to the fluorescence intensity identifies with sub-pixel accuracy the centroid of the fluorescence, and therefore of the ion channel(s) producing the Ca^2+^ signal. Top panel shows a representative images of a region of interest showing a processed, filtered Piezo1 flicker event (red arrow). Middle panel shows a 3D representation of the event and the bottom panel shows a Gaussian fit to the event. Arrow in bottom panel marks the peak of the Gaussian profile that identifies with sub-pixel accuracy the location of the centroid of the Ca^2+^ flicker. Scale bar = 10 µm. **D.** Flicker localizations (red dots) overlaid on an image of a MEF to map sites of Piezo1 activity. **E.** Flicker activity from the region of interest marked in D plotted over time as fluorescence ratio changes (ΔF/F_0_). Identified flicker events are highlighted in red. Scale bars = 10 µm. See also Movie M3 and Fig. S2.

Figure 2 shows an implementation of the algorithm applied to Piezo1 Ca^2+^ flickers recorded from hNSPCs (see also Movie M3 and Fig. S2). Piezo1 Ca^2+^ flickers are visualized by imaging Ca^2+^ influx through the channel using TIRFM (Fig. 2A). The raw movie is processed to produce a F/F_0_ ratio movie (Fig. 2B) which is then spatially and temporally filtered to increase the signal-to-noise ratio of the signals of interest. The processed movie is passed through the clustering algorithm for event detection. Once events are detected, a 2-dimensional (2D) Gaussian curve is fit to every event in the movie to determine the localization of each flicker event with subpixel precision. Figure 2C shows the output of the algorithm for a single, representative flicker event after pre-processing steps (Fig. 2C, top and middle) and after the subpixel localization of the event by Gaussian fitting (Fig. 2C, bottom). The peak of this 2D Gaussian (red arrow, Fig. 2C bottom) identifies the center of the Ca^2+^ event with subpixel accuracy. Assuming the diffusion of Ca^2+^ is radially symmetric, this gives the location of an individual ion channel, or the ‘center of mass’ of the group of ion channels, underlying the event. These flicker localizations are overlaid on an image of the cells (Fig. 2D) to produce a cellular map of active Piezo1 channels. The extracted signals can be analyzed to determine peak amplitude, temporal dynamics, and frequency of signals at a specific site (Fig. 2E). This technique made it possible for us to examine the spatial localization of Piezo1 activity in relation to cellular traction forces.

### Piezo1 Ca^2+^ flickers are enriched at regions predicted to have high traction forces

To relate spatial maps of Piezo1 Ca^2+^ flicker activity to cellular traction forces, we mapped Piezo1 activity in cells with known patterns of traction forces. We utilized the well-established effect of cell geometry on traction forces: cell shape determines *where* forces are generated and cell size determines *how much* force is generated ^39–42^. We controlled the shape and size of HFFs and hNSPCs -- and therefore the spatial pattern and magnitude of their cellular traction forces -- using substrate micropatterning ^42–44^ and examined Piezo1 Ca^2+^ flicker maps in these micropatterned cells. To do so, glass coverslips were patterned with islands of fibronectin of pre-determined shapes and sizes. Upon seeding, cells bind to fibronectin via cellular integrins and take up the geometry of the island. We selected the shape of our substrate islands based on previous traction force measurements in micropatterned cells ^39–41,45^, which show that in cells constrained to a square shape, traction forces are highest at the vertices, moderately high at edges, and minimal in the middle of the cell (Fig. 3A). Moreover, as the size of the square island is increased, the magnitude of traction force increases ^40,42^. This robust dependence of traction forces on the shape and size of micropatterned square cells allowed us to ask whether the location and magnitude of Piezo1 Ca^2+^ flickers in square cells also show a similar dependence on cell shape and size.

**Figure 3.**
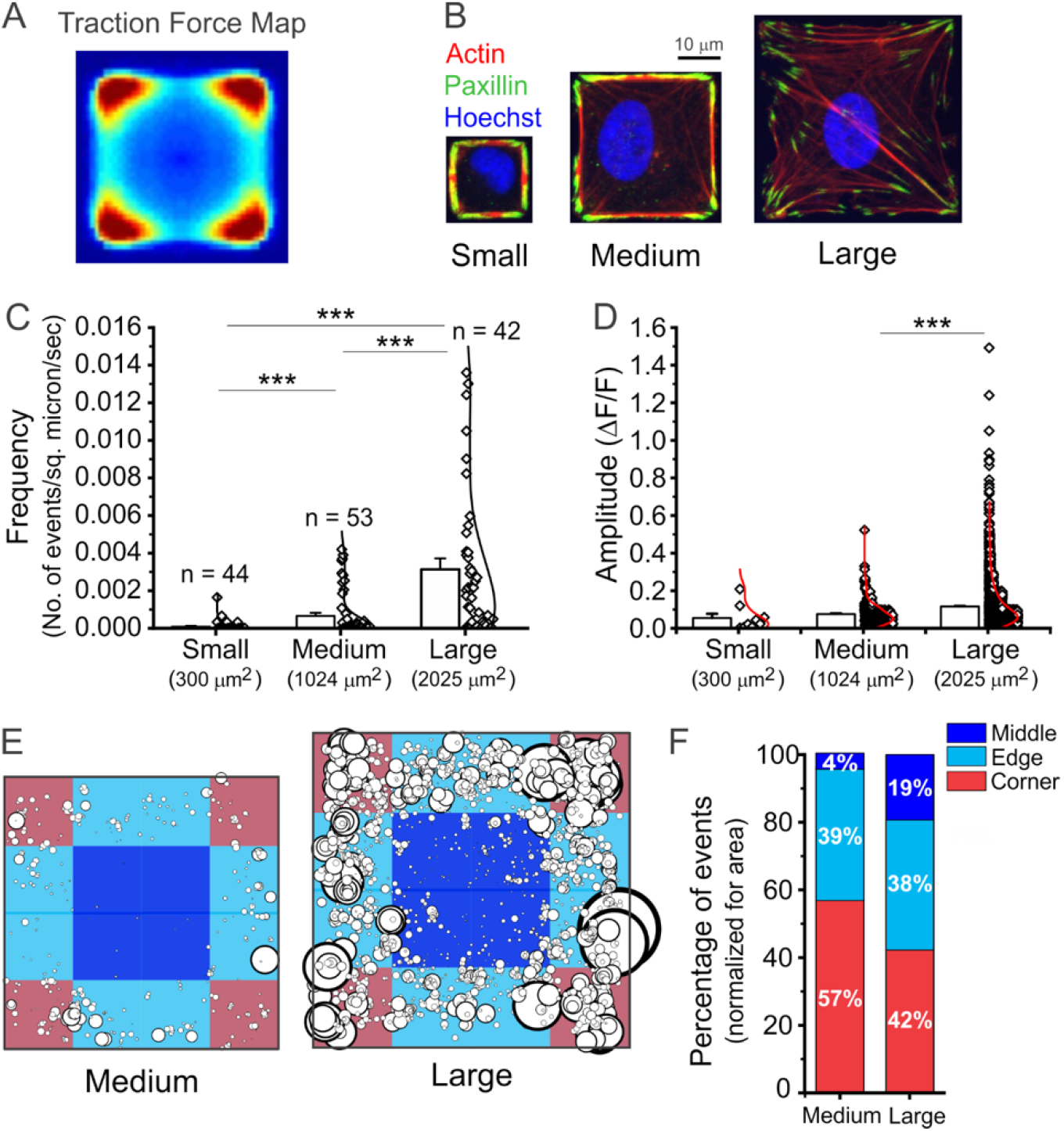
Piezo1 Ca^2+^ flickers are enriched in regions of cells predicted to have higher traction forces. **A.** Cells constrained to adopt a square shape on a patterned substrate generate the largest traction forces at the corners (red), lower levels of forces at the edges (cyan) and lowest traction forces in the middle of the cell. The image is reproduced from Holt et al. 2012 under CC BY-NC-SA 3.0 license. **B.** Cells seeded on square fibronectin islands yield a single square cell per island. Images are representative confocal slices of hNPSCs stained for the actin cytoskeleton (phalloidin, red), the focal adhesion zone protein, paxillin (anti-paxillin antibody, green), and the nucleus (Hoechst, blue). Note the larger number of actin stress fibers terminating in focal adhesions as cell spread area increases. Island sizes used: Small, 300 µm^2^; Medium 1024 µm^2^; Large, 2025 µm^2^. **C.** Cells on larger square islands display more Piezo1 Ca^2+^ flickers as evidenced from the frequency of Piezo1 Ca^2+^ flicker events for HFF cells seeded on Small, Medium, and Large islands. Number of cells imaged for each size is represented in the graph. *** p < 0.001 by Kolmogorov-Smirnov test. **D.** Flicker amplitudes are larger in cells with larger spread area. Data are from 9 flicker events from 44 Small cells, 355 events from 53 Medium and 2411 events from 42 Large cells. *** p < 0.001 by Kolmogorov-Smirnov test. **E.** Localization and amplitude of flicker events in Medium and Large cells. Location of each flicker events from Medium and Large cells is represented on a square in which the Corner (red), Edge (Cyan) and Middle (Blue) regions are marked. Each circle represents the site of a flicker event, with the size of the circle scaled by the amplitude of the flicker. **F.** Quantitation of flicker events from panel E in Small, Medium and Large areas of square islands. A chi-square test was performed to determine whether the observed distribution is different from chance: for Medium cells χ^2^(2, N = 355) = 108.38, p < 0.0001 and for Large cells, χ^2^(2, N = 2411) = 161.89, p < .0001. See also Fig. S3.

We seeded cells on glass substrates in square shapes of three different sizes (Small 17.3 µm × 17.3 µm= 300 µm^2^, Medium 32 µm × 32 µm = 1024 µm^2^, Large 45 µm × 45 µm = 2025 µm^2^). We confirmed that micropatterned cells exhibited the shape and cytoskeletal organization expected of this geometry. For this, we visualized actin filaments in fixed micropatterned cells with fluorescently-labeled phalloidin, focal adhesions with an anti-Paxillin antibody, and cell nuclei with Hoechst dye (Fig. 3B). Cells on larger islands displayed greater numbers of, and longer, actin stress fibers, terminating in paxillin-rich focal adhesions that were concentrated in corner regions. Cells on Large islands displayed a network of actin stress fibers across the cell, while cells on Small islands showed actin accumulated primarily along the edges, as previously observed in other cell types for this specific set of square patterns ^46^.

We next imaged Piezo1 Ca^2+^ flickers in live cells adhering to Small, Medium, and Large islands. Flicker activity was observed in 6 out of 44 (13.6%) Small cells, 29 out of 53 (54.7%) Medium cells and 38 out of 42 (90.5%) Large cells. Quantification of flicker frequency from all cells imaged additionally showed that flicker frequency scaled with cell size (Fig. 3C). The amplitudes of Piezo1 Ca^2+^ flickers also differed in Small, Medium and Large cells, with larger cells which are known to generate larger traction forces displaying larger flickers (Fig. 3D). To determine the location of Piezo1 Ca^2+^ flicker activity relative to the predicted traction force distribution, we examined flicker localizations from Medium and Large cells; Small cells were not included in this analysis due to the small number of flicker events observed. We determined the localization of flickers for Medium and Large cells in three regions and found that Corner and Edge regions showed a higher number of flicker events, which were also larger in amplitude than events in the Middle region (Fig. 3E). If flickers were evenly distributed, we would expect an equal occurrence in Corner, Middle, and Edge regions once normalized for area. However, we observed that Corner regions showed more flickers, followed by Edge regions and Middle regions (Fig. 3F). Similar results were also observed for hNSPCs (Fig. S3). Overall, our measurements show that Piezo1 Ca^2+^ flickers are enriched in regions of the cell expected to have higher traction forces, and that Piezo1 Ca^2+^ flicker frequency and amplitude scales with cell spread area.

### Piezo1 Ca^2+^ flickers localize to hotspots of traction forces

Our finding of enriched Piezo1 Ca^2+^ flickers in regions of micropatterned hNSPCs predicted to have higher traction forces motivated us to measure traction forces and Piezo1 Ca^2+^ flickers in the same cell. We used a Förster resonance energy transfer (FRET)-based molecular tension sensor (MTS) to measure cellular traction forces ^47^. The MTS is comprised of an elastic peptide which separates a covalently-bound FRET pair (Fig. 4A). The N-terminus of the sensor is covalently attached to a functionalized glass coverslip to produce a carpet of sensors. The C-terminus of the MTS has a fibronectin fragment to which cells bind via integrins, such that cells seeded on to the sensor-coated glass coverslip adhere to the substrate via the MTS. Traction forces generated by the cell are communicated to the MTS via integrin-MTS attachments: these forces cause extension of the peptide spring, leading to a separation of the FRET pair and therefore a reduction in FRET efficiency. The FRET donor and acceptor channels are simultaneously imaged, yielding FRET index maps that are calculated by dividing the acceptor intensity over the sum of donor and acceptor intensities. A high FRET index indicates low traction force and a low FRET index indicates high traction force. The FRET index maps can be converted to FRET efficiency maps to allow extraction of quantitative values for force based on the FRET-force response of the elastic peptide ^48,49^; see Methods for details). Thus, imaging the donor and acceptor fluorophores allows the production of a quantitative, high-resolution traction force map of the cell. Compatibility of these sensors with TIRFM-based Ca^2+^ imaging allowed us to measure and correlate cellular traction forces and Piezo1 activity in the same cell.

**Figure 4.**
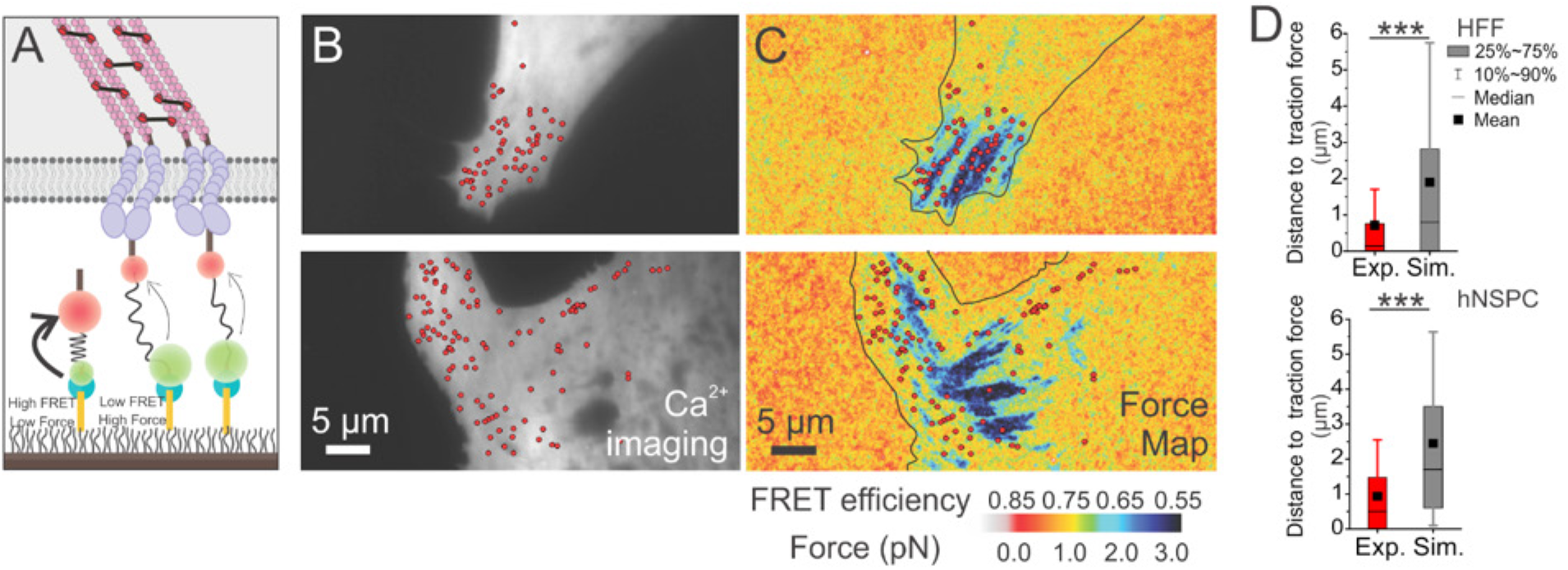
Piezo1 Ca^2+^ flickers localize to regions of high traction forces. **A.** Schematic of the Molecular Tension Sensor (MTS) used for traction force imaging. The N-terminal region of the sensor (blue) is tethered to a PEG-functionalized glass coverslip. A fibronectin domain at the C-terminal end (brown) binds to the cell’s integrins (purple), allowing cells to attach to the glass coverslip. An elastic spring domain bridges the two ends of the sensor and separates a FRET donor (green) and acceptor (red). Cell-generated traction forces pull the FRET pair apart, resulting in reduced FRET efficiency. The FRET index (the ratio of acceptor intensity over summed donor and acceptor intensities), serves as a measure of force; with a low FRET index indicating high force, and a high FRET index indicating a low force. **B.** Imaging of Piezo1 Ca^2+^ flickers. Panels show resting fluorescence of HFF cells loaded with Ca^2+^ indicator Cal-520, with overlaid red dots marking the centroid locations of Ca^2+^ flickers. **C.** Corresponding force maps from the same cells, overlaid with red dots marking the Ca^2+^ flicker locations. Blue denotes low FRET (high force) and red denotes high FRET (low force). The color bar in C represents FRET efficiency (top) and the average force per MTS per pixel in pN obtained from calibrated FRET-Force curves of the MTS as described in Methods. **D.** Box and whisker plots with red boxes showing distances from Piezo1 flicker localizations to the nearest traction force region for HFFs (top: 515 flickers from 9 cells) and hNSPCs (bottom: 66 flickers from 18 cells). Grey boxes show corresponding mean distances derived from simulations of 9000 random intracellular locations for each cell. Box range is 25^th^ to 75^th^ percentile; whiskers denote 10^th^ and 90^th^ percentile, horizontal lines represent the median and filled black squares represent mean. *** denotes p < 0.001 by Kolmogorov-Smirnov test.

We imaged force maps and Ca^2+^ flickers in HFFs, a popular cell type for studying traction forces because they display large adhesions which generate high traction forces ^47,48,50^. We seeded HFFs onto coverslips functionalized with the MTS, allowed the cells to attach and spread for 1-2 hours, then loaded them with the Ca^2+^ indicator Cal-520 AM. We imaged traction forces followed by Piezo1 activity (Fig. 4B, C). Overlaying maps of Piezo1 Ca^2+^ flickers and force demonstrated that Piezo1 Ca2+ flickers occurred in regions of the cell that displayed high traction forces (Fig. 4C). To quantify the spatial relationship between traction forces and Piezo1 Ca^2+^ flickers, we calculated the distance of Piezo1 Ca^2+^ flickers to the nearest force-producing region (Fig. 4D). To determine whether the localization of Piezo1 Ca^2+^ flickers was different from chance, we simulated 9000 randomly localized Piezo1 Ca^2+^ flicker sites in each cell and compared the distance of experimental and randomly simulated Piezo1 Ca^2+^ flicker localizations to the nearest high-force region. On average, flicker localizations were located 0.72 µm from force-producing adhesions, whereas simulated flicker localizations were located at a distance of 1.9 µm from force-producing regions (Fig. 4D, top; p < 0.001 by Kolmogorov-Smirnov test). In similar experiments with hNSPCs we found that experimental flickers were 0.94 µm away from high-force regions, whereas simulated flicker localizations were situated 2.2 µm away (Fig. 4D, bottom; p <0.001 by Kolmogorov-Smirnov test). Together, our findings indicate that Piezo1 Ca^2+^ flicker location is spatially correlated with higher traction forces.

### Piezo1 channels diffuse over the surface of the cell

The traction force produced by Myosin II is communicated through actin filaments to focal adhesions that attach to the substrate ^32,51^. Our observation that Piezo1 Ca^2+^ flickers arise predominantly in the vicinity of force-producing focal adhesions suggested two possibilities: (i) Piezo1 channels are localized to focal adhesions where traction forces are transmitted to the substrate, or (ii) Piezo1 channels are present all over the cell surface, but are only activated by traction forces near force-producing adhesions. To distinguish between these possibilities we visualized the localization of Piezo1 proteins. The dearth of sensitive and specific antibodies against endogenous Piezo1 precluded immunolocalization of the native channel to answer this question. Instead, we used a knock-in reporter mouse wherein a tdTomato fluorescent protein is tagged to the C-terminus of the endogenous Piezo1 channel ^5^ (Fig. S4A). The expression of the Piezo1-tdTomato fusion protein is driven by the native Piezo1 promoter and regulatory elements; thus expression levels and patterns of the tagged channel are expected to be the same as that of endogenous channels. We immunostained endogenous Piezo1-tdTomato channels in mNSPCs with an anti-RFP antibody and observed channels distributed all over the cell surface rather than being localized to focal adhesions (Fig. S4B, C). Imaging of the tdTomato moiety in live mNSPCs at the cell-substrate interface by TIRF microscopy revealed channel localization over the ventral surface of the cell (Fig. 5A) and that individual Piezo1 puncta are mobile in the plasma membrane (Movie M4). We tracked mobile tdTomato-tagged Piezo1 channel puncta in the plasma membrane in images captured every 100 ms with TIRFM using custom-written single particle tracking scripts (See Methods) to build trajectories of individual Piezo1 puncta (Fig. 5B). Fig. 5C shows the atracks of 10 randomly chosen trajectories in a ‘flower plot’. To obtain apparent diffusion coefficients, we plotted the ensemble Mean Squared Displacement, MSD, of 5,965 tracks. The slope of the MSD yields an apparent two-dimensional diffusion coefficient of 0.067µm^2^/s, which is similar to that of other membrane proteins ^52–54^.

**Figure 5.**
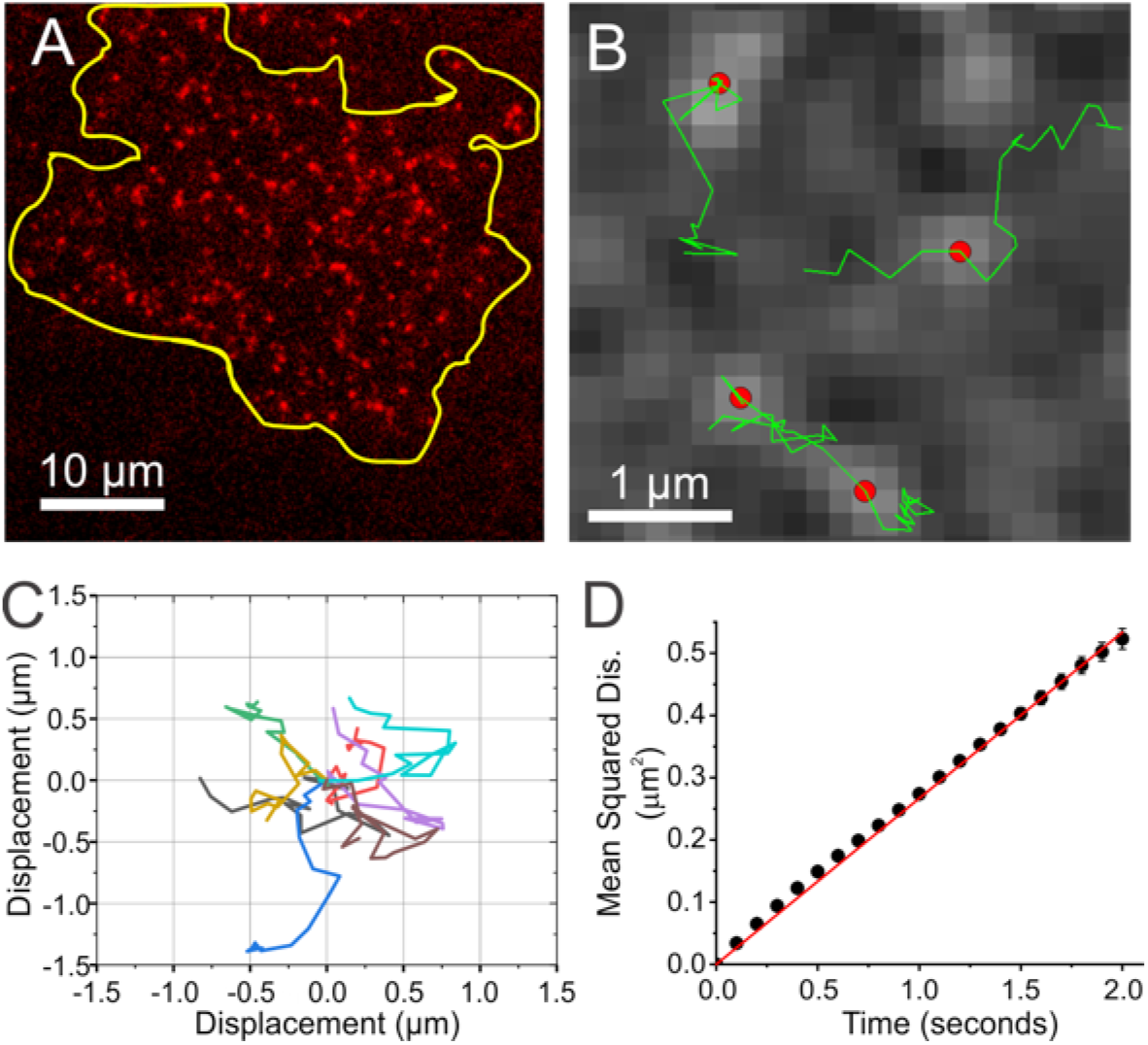
Piezo1 molecules are mobile in the plasma membrane. **A.** Representative TIRF image of tdTomato fluorescence from the cell-substrate interface of live mNSPCs harvested from Piezo1-tdTomato reporter mice. Yellow line denotes the cell boundary of the cell. Puncta visible outside the yellow cell line are from neighboring cells. Note the widespread distribution of Piezo1-tdTomato channels over the cell surface. **B.** Tracking of Piezo1-tdTomato channel puncta imaged at 10 frames per second in live mNSPCs reveals motility of channel puncta. The background image shows fluorescence of Piezo1-tdTomato puncta captured during a single imaging frame. Red dots mark the localizations of the puncta during that frame, and green lines depict the tracks of these puncta over several successive frames. **C.** ‘Flower plot’ derived by overlaying the trajectories of 10 individual puncta over 1 second after normalizing their starting coordinates to the origin. **D.** Mean-squared displacement calculated from 6853 Piezo1-tdTomato tracks plotted as a function of time. Error bars represent standard error of the mean. Error bars are smaller than data symbols for most points. The data fit to a straight line with a slope corresponding to a 2D diffusion coefficient of 0.067 µm^2^/s. R^2^ for linear fit to data is 0.99. See also Fig. S4 and Movie M4.

Taken together, the widespread distribution of Piezo1 channels on the ventral surface of the cell and their mobility suggest that channels in the vicinity of focal adhesions are activated by local mechanical stresses produced by traction force generation, whereas the channels farther away remain largely silent.

### Force generation by MLCK-mediated Myosin II phosphorylation activates Piezo1

Nonmuscle Myosin II hydrolyzes ATP to convert chemical energy into mechanical force, which is communicated through actin filaments and focal adhesions to the extracellular matrix (Fig. 6A). We previously showed that inhibition of Myosin II by blebbistatin inhibited Piezo1 Ca^2+^ flickers ^7^, establishing that force generation by Myosin II is required for Piezo1 Ca2+ flicker activity. Myosin II activity is regulated by the Myosin II regulatory light chain subunit, whose phosphorylation converts Myosin II from an inactive form to an active form capable of filament assembly and force generation (Fig. 6A). We asked how the phosphorylation state of Myosin II might impact Piezo1 activity. Myosin II is phosphorylated by two kinases - Rho-associated protein kinase (ROCK) and Myosin Light Chain Kinase (MLCK). The two kinases control distinct spatial pools of Myosin II: ROCK phos-phorylates Myosin II in the center of the cells while MLCK phosphorylates Myosin II in the periphery ^55–57^. The ROCK inhibitor Y-27632 had no effect on Piezo1 Ca2+ flicker frequency (Fig. 6B). On the other hand, the MLCK inhibitor ML-7, which we previously showed to rapidly reduce traction force generation in HFFs ^48^, effectively inhibited Piezo1 Ca^2+^ flickers (Fig. 6C). The regulation of Piezo1 Ca^2+^ flickers by MLCK (which has been shown to phosphorylate Myosin II at the periphery of the cell) but not by ROCK (which activates Myosin II in the center of the cell) is consistent with our observation that Piezo1 Ca^2+^ flickers are more often observed in the periphery of cells (for example, see Piezo1 activity maps in Figs. 3 and 4).

**Figure 6.**
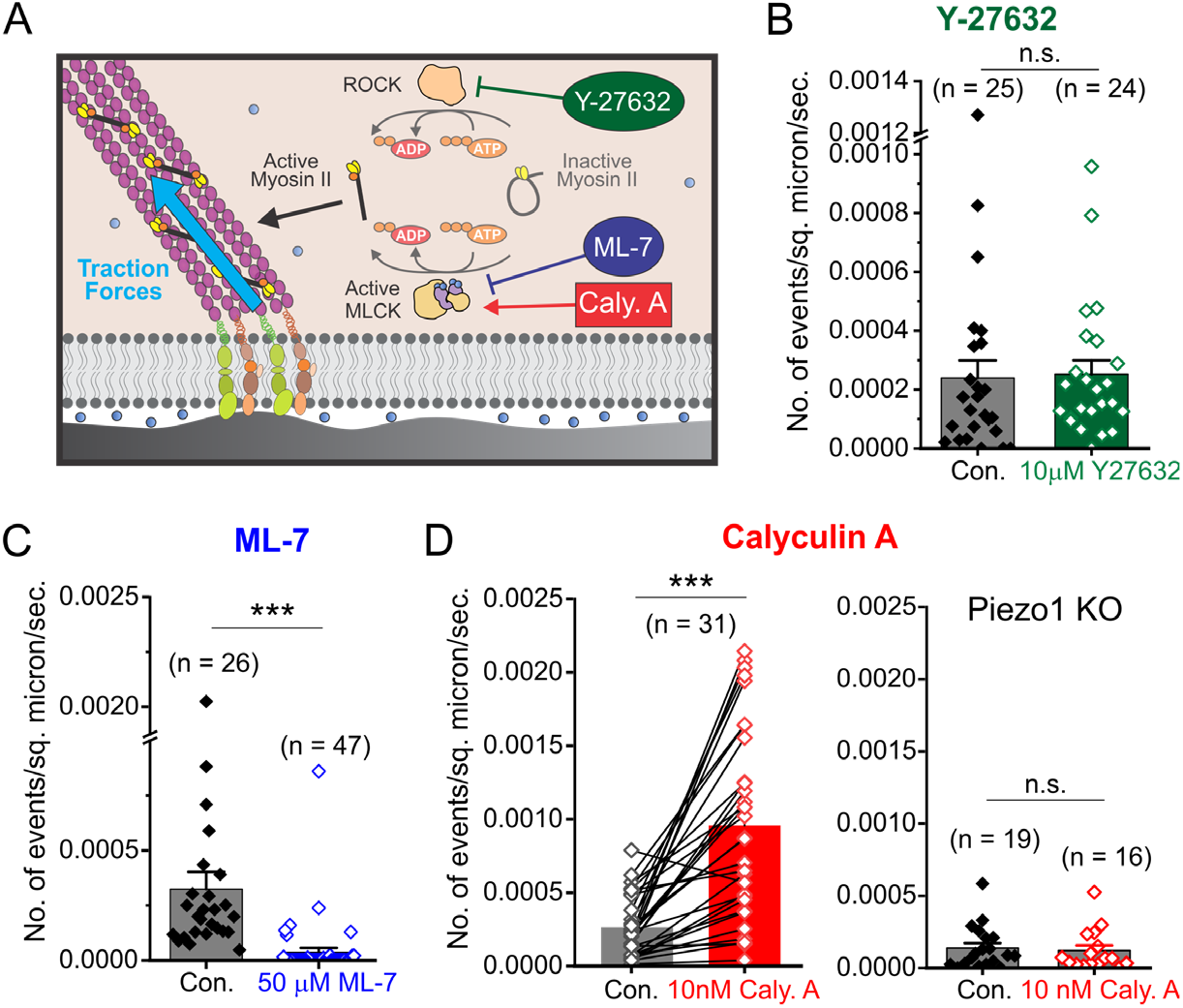
Piezo1 Ca^2+^ flickers are evoked by MLCK activity. **A.** Schematic of traction force generation by Myosin II. The enzymes Rho-associated Protein Kinase (ROCK) and Myosin Light Chain Kinase (MLCK, beige and purple) phosphorylate Myosin II (yellow and black) to generate the activated form that binds actin filaments (purple) and produces traction forces (blue arrow). The drug Y-27632 inhibits ROCK, the drug ML-7 inhibits MLCK and the drug Calyculin A potentiates MLCK activity, to alter traction force generation by Myosin II. **B.** Bar and data plot of Piezo1 flicker frequency from WT HFFs in Control imaging solution and in the presence of 10 µM ROCK inhibitor Y-27632 showing that the drug has no effect on Piezo1 Ca^2+^ flickers frequency. **C.** Treatment of HFFs with 50 µM ML-7 inhibits Piezo1 Ca^2+^ flickers. **D.** Treatment of HFFs by 10 nM Calyculin A increases the frequency of Piezo1 Ca^2+^ flickers. Left, Paired measurements of flicker frequency in the same fields of view in Control imaging solution (gray) and 1-5 minutes after replacement of imaging solution with solution containing 10 nM Calyculin A (red). Right, 10 nM Calyculin A does not increase Ca^2+^ flickers in Piezo1-Knockout HFFs (n.s. denotes “not statistically significant” and *** denotes p < 0.001 by Kolmogorov-Smirnov test). In each panel, data are from three experiments; bar height denotes the mean of the data points, error bars denote standard error of the mean, and each point represents mean flicker frequency in an individual video; the number of videos analyzed from 3 different experiments is specified above each bar. See also Fig. S5.

Previous work establishes that treatment of cells with Calyculin A, an inhibitor of myosin light chain phosphatase, increases myosin II-dependent force generation ^58–60^. We found that Ca^2+^ flickers in the same set of HFFs before and after treatment with 10 nM Calyculin A showed on average of 5-fold increase in Ca^2+^ flickers within minutes (Fig. 6D, left). Calyculin A failed to increase flicker activity in the absence of external Ca^2+^, indicating that Ca^2+^ influx across the plasma membrane is required (Fig. S5). Piezo1 KO HFFs did not show increased flicker activity in response to Calyculin A (Fig. 6D, right), indicating that the observed increase in frequency of Ca^2+^ flickers is mediated by Piezo1. In summary, we demonstrate that traction forces produced by nonmuscle Myosin II induce spatially restricted Ca^2+^ flickers by activating Piezo1 channels, and identify MLCK-mediated phosphorylation of Myosin II as an upstream signaling mechanism that regulates the force generation.

## DISCUSSION

### Technical advances in Piezo1 activity measurements

Emerging evidence for a functional interplay of Piezo1 and the cellular cytoskeleton ^1^ emphasizes the need for studying Piezo1 activity in native cellular conditions and in conjunction with cytoskeletal dynamics. To measure the activity of native Piezo1 channels in intact cells, we use a TIRFM-based assay of Piezo1 Ca^2+^ flickers to monitor Piezo1 activity with millisecond temporal and sub-micron spatial resolution. The high signal-to-noise ratio afforded by TIRFM allowed detection of small signals arising from the activity of endogenously-expressed channels at the cell-substrate interface. We developed a custom-written, open-source analysis algorithm that utilizes principles from localization microscopy for the automated detection, localization, and measurement of Ca^2+^ flickers. This approach enabled us to generate overlaid spatial maps of Piezo1 Ca^2+^ flickers and cell-generated traction forces in the same cell. Thus, we provide a novel experimental and analytical framework for examining the interplay between Piezo1 and the cytoskeleton in the native cellular environment.

We employ these technical advances to demonstrate the presence of local, discrete Ca^2+^ flickers at the cell-substrate interface that are dependent on Piezo1 expression and are elicited in a spatially-restricted manner that requires Myosin II activation through MLCK. The marked reduction of Ca^2+^ flickers in Piezo1-deficient cells (Fig. 1, S1 and ref. ^7^) and in the absence of extracellular Ca^2+^ (Fig. S1 and ref. ^7^) demonstrate that under the experimental conditions of this study, Ca^2+^ flickers are generated primarily through the action of plasma membrane-localized Piezo1. Together with our previous work ^7^, our findings constitute a novel mode of activating the Piezo1 channel that may be relevant in a variety of physiological contexts. Our study also provides mechanistic insights for how spatially-localized Ca^2+^ flickers through ion channels may be elicited in response to traction forces. We show that the spatial restriction of Ca^2+^ flickers doesn’t arise from localized expression of channels to focal adhesions; rather, channels are mobile and localized flicker activity is generated by selective activation of channels near force-producing focal adhesions. A similar mechanism may also apply to some Trp channels (e.g. TrpM7 and TrpC1) for which flicker activity in the vicinity of focal adhesions or preferentially on stiff substrates has been reported ^61,62^.

Our approach complements electrophysiological assays of Piezo1 activity. Most studies of Piezo1 activation have utilized patch clamp recording of ionic currents through the channels. In whole-cell patch clamp, cellular contents are dialyzed by the large reservoir of solution in the patch pipette, confounding the study of channel activation and modulation by the cytoskeleton and by soluble intracellular molecules. In cell-attached patch clamp, the intracellular contents are retained, but the gigaseal connection between the membrane and glass pipette exerts intense mechanical stress on the membrane patch ^65^. This is sufficient to drive a large fraction of Piezo1 channels into inactivation ^36^, resulting in a higher activation threshold compared to physiological conditions. In comparison, our assay does not disrupt the cellular cytoskeleton or dialyze the cell, providing a measurement of channel dynamics under native cellular conditions, and allowing spatial monitoring of sub-cellular domains of Piezo1 activity that is not feasible with patch clamp electrophysiology.

### Spatial regulation of Piezo1 activity by cellular traction forces

We combined our Piezo1 Ca^2+^ flicker assay with approaches to manipulate and measure intrinsic cellular traction forces. First, we used micropatterned square substrates to constrain the shape and size of cells such that they generate a known pattern of traction forces ^42–44^. Piezo1 Ca^2+^ flickers were enhanced in corners and edges of these cells - regions predicted to have high traction forces (Fig. 3). Second, we used a FRET-based molecular tension sensor (MTS) ^47,50^ to spatially-resolve and quantitatively measure cellular traction forces, that we correlated with Piezo1 activity in the same cell. These measurements would be difficult using conventional traction force microscopy (TFM), which tracks the displacement of fluorescent beads in a soft gel substrate, due to the technical challenges inherent in imaging Ca^2+^ flickers on soft substrates, as well as the limited spatial resolution of commonly implemented versions of TFM. We observed a clear spatial correspondence between Piezo1 Ca^2+^ flickers and high traction forces, consistent with local cellular traction forces activating the channel. Moreover, we elucidate an upstream signaling mechanism involving phosphorylation of Myosin II by MLCK as responsible for the generation of the force that activates Piezo1. Conversely, the Myosin II kinase ROCK does not seem to be involved in generating Piezo1 Ca^2+^ flickers. Given that MLCK is itself regulated by Ca^2+^, we speculate that MLCK, nonmuscle myosin II, and Piezo1 might constitute a feedforward loop whose activity may enhance myosin contractility in regions of the cytoskeleton proximal to load-bearing attachments to the ECM. Moreover, little is known of how cells detect the traction forces that they themselves generate. We propose that Piezo1 plays an important role in that regard, and provide evidence that it’s activity is localized. The local nature of Piezo1 Ca^2+^ flickers in turn suggests that they may locally regulate contractility. Interestingly, Ca^2+^ influx through unidentified stretch-activated ion channels was previously shown to precede an increase in traction forces ^63,64^. The complex relationship between Ca^2+^ influx and traction forces opens the possibility of a feedback loop in which traction forces activating Piezo1 become stronger as a result of Piezo1-dependent calcium signaling. It also allows for cross-talk between other sources of calcium influx and Piezo1 activity. These interesting possibilities warrant further investigation.

### Motility and clustering of Piezo1 channels

We find that Piezo1 channels are mobile in the cell membrane, with an apparent ensemble diffusion coefficient of 0.067 µm^2^/s. This value is within the wide range of diffusion coefficients of 0.01 - 0.3 µm^2^/s measured for membrane proteins ^52–54,66^. Whereas Piezo1 channels appear to diffuse readily in the plasma membrane, the restriction of flicker activity to regions of the cell that exhibit traction forces (Fig. 4) raises the possibility that active channels may be transiently anchored. A full analysis of the subcellular localization dynamics of Piezo1 is beyond the scope of this study, but is likely to provide key insights into Piezo1-mediated mechanotransduction and the interaction of the channel with its cellular environment.

An open question is whether Piezo1 Ca^2+^ flickers represent the activity of single channels or a cluster of channels, and correspondingly, whether the motile Piezo1-tdTomato puncta represent individual channels or clusters of channels that move as a unit, as has been described for IP_3_ receptors ^67^. We observed a larger amplitude of Piezo1 Ca^2+^ flickers in larger cells, which have higher traction forces (Fig. 3). If flickers represent single-channel activations, then we would expect to observe changes in flicker frequency but not in amplitude. Thus, it is plausible that flickers represent the activity of clusters of channels, with higher forces activating a larger fraction of channels in the cluster. Consistent with this idea, Bae et al. ^21^ observed in cell-attached patch clamp experiments that groups of Piezo1 channels sometimes showed a collective change in dynamics, including a collective loss of inactivation or an abrupt change in activation kinetics. Alternatively, the measured amplitude differences could arise from bursts of unresolved individual openings.

### How do traction forces activate Piezo1?

Several studies have proposed that Piezo1 is gated by membrane tension ^27,28,36,68^, and three recent cryo-EM structures of Piezo1 ^68–70^ support this gating mechanism. We sometimes observed Piezo1 Ca^2+^ flickers located a few microns proximal to, but not directly overlying the traction force hotspots (Fig. 4C). The parsimonious explanation for this observation is that mechanical stress may be communicated to the channel through the plasma membrane. Our working model for the activation of Piezo1 Ca^2+^ flickers by traction forces is that traction forces produce a local increase in membrane tension that activates Piezo1 channels in the vicinity of force-producing adhesions (Fig. 7).

**Figure 7.**
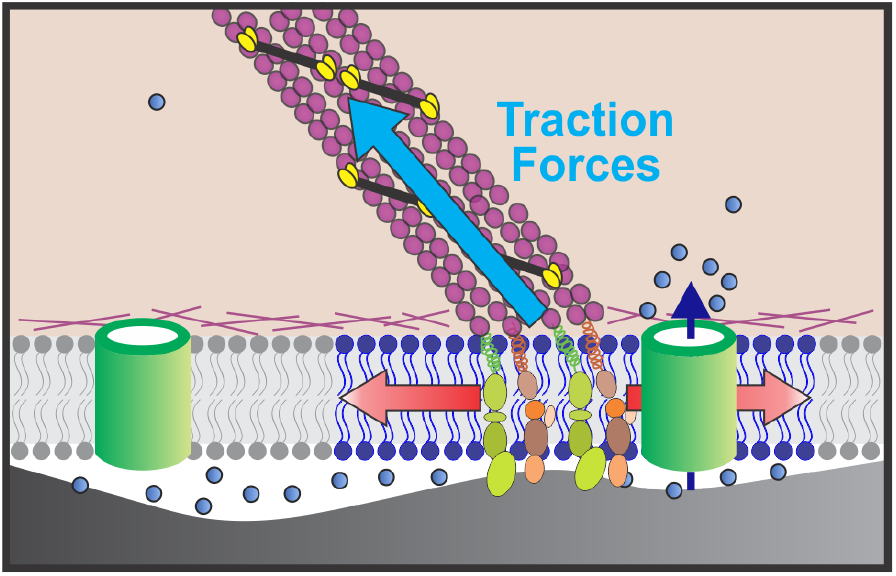
Working model of Piezo1 activation by traction forces. Traction forces (blue arrow) generated by Myosin II motors (yellow and black) along actin filaments (purple) tethered to integrin-based focal adhesion zones (green and brown) generate local increases in membrane tension (depicted by blue region of lipid bilayer and red arrows) that elicit Ca^2+^ flickers from nearby Piezo1 channels (green cylinder in right part of figure). Blue dots represent Ca^2+^ ions and dark blue arrow represents Ca^2+^ influx through Piezo1. Piezo1 channels far from force-producing adhesions are not activated (green cylinder in left of figure).

Whether membrane tension is a global or a local cellular parameter has been a subject of ongoing debate ^71^. A recent study demonstrates that in intact cells -- unlike in membrane blebs -- perturbation of membrane tension can be a local event that does not necessarily propagate beyond a few microns ^72^, a finding that is supported by the activation of the bacterial mechanosensitive channel MscL in mammalian cells ^73^. Our model that local membrane tension induced by cytoskeletal forces activates Piezo1 is consistent with these reports. However, we cannot presently exclude contributions from transient physical interactions between Piezo1 and focal adhesion proteins, or from changes in membrane organization that may occur near traction force regions.

### An emerging picture of Piezo1 mechanotransduction

Piezo1 responds on the millisecond timescale to diverse external mechanical cues such as cell indentation ^2^, shear flow ^4^, membrane stretching ^2,36^, substrate displacement ^30^, and osmotic stress ^28^. Some of these mechanical stimuli impinge upon a small region of the cell, whereas others affect the cell in its entirety. How may Piezo1 channels respond to mechanical cues that may strike anywhere and at any time in the cell while also transducing cell-generated traction forces that occur specifically at focal adhesion zones? We propose that -- like policemen patrolling a city -- mobility allows a smaller number of Piezo1 channels to explore a larger number of mechanical microdomains, and thereby respond to a greater diversity of mechanical cues. For instance, recurrent local mechanical stimuli may be entirely missed by sparsely distributed, static channels; however mobility would allow channels to detect at least a subset of the events. Whereas the electrical signal generated from Piezo1 ion flux would globally depolarize the cell, the restricted nature of Ca^2+^ diffusion in the cytosol tightly constrains the ‘chemical’ signal to the vicinity of the channel. Thus, spatial localization of Piezo1 activity could serve to spatially localize biochemical signaling downstream of Piezo1, and may be a key component rendering specificity to its diverse physiologic roles in different cell types.

## ACKNOWLEDGEMENTS

We thank Truc Tran, Nhu Nguyen, Huixun Du, Klara Zakery, Juhi Gopal, Brian Nguyen, and Chang Zhao for technical assistance; Vivian Leung for help with manuscript preparation; Dr. Douglas Tobias and members of Pathak lab for discussions; Dr. Ardem Patapoutian for the gift of Piezo1-tdTomato mice; and the University of California Natural Reserve System (Steele/Burnard Anza-Borrego Desert Research Center Reserve DOI: 10.21973/N3Q94F) for providing a venue for scientific discussions. This work was supported by NIH grants DP2 AT010376 and R01 NS109810 to M. M. P., R21 NS085628 to F. T., R37 GM048071 to I.P, F31 GM119330 to K.E., R01 GM112998 and an HHMI Faculty Scholar Award to A.R.D., CIRM RB5-07254 to L.A.F. and T32 NS082174 to J.A.

## Materials and Methods

### Cell Culture

#### hNSPC Culture

All research involving human cells was approved by the University of California, Irvine Institutional Review Board and the Human Stem Cell Research Oversight Committee, and had no patient identifiers. Brain-derived fetal hNSPC cultures (SC27) were isolated from the cerebral cortex of a male fetus of 23-wk gestational age and were maintained as previously described (Pathak et al. 2014). Briefly, undifferentiated cells were grown as adherent cultures on fibronectin (Fisher Scientific)-coated flasks in basal medium containing DMEM/F12 (GIBCO), 20% BIT-9500 (Stem Cell Technologies), and 1% antibiotic/antimycotic (Invitrogen) supplemented with the following growth factors: 40 ng/mL EGF (BD Biosciences), 40 ng/mL FGF (BD Biosciences), and 40 ng/mL PDGF (Peprotech). hNSPCs were passaged approximately every 5-7 days using Cell Dissociation Buffer (Invitrogen) and split 1:2. Cells were used at passages P10–22. Informed written consent was obtained for all human subjects.

#### mNSPC Culture

All studies were approved by the Institutional Animal Care and Use Committee at UCI. NSPCs from cerebral cortices of E12.5 wildtype mice or from mice expressing a C-terminal fusion of Piezo1 with tdTomato (Piezo-tdTomato) (Ranade et al. 2014) were cultured as neurospheres as described previously (Nourse et al. 2014). Piezo1-tdTomato reporter mice were a gift from A. Patapoutian. mNSPC growth medium consisted of: High glucose Dulbecco’s modified Eagle’s medium (all reagents from Life Technologies unless otherwise noted), 1× B27, 1× N2, 1 mM sodium pyruvate, 2 mM glutamine, 1 mM N-acetylcysteine (Sigma Aldrich), 20 ng/ml epidermal growth factor (EGF) (BD Biosciences), 10 ng/ml fibroblast growth factor (FGF) (BD Biosciences), and 2 µg/ml heparin (Sigma Aldrich). Cells were passaged by dissociation with Neurocult Chemical Dissociation Kit (Stem Cell Technologies). For immunostaining, NSPCs and HFF cells were plated on #1.5 glass coverslips (Warner Instruments). For live cell TIRFM imaging, mNSPCs cells were plated on #1.5 glass Mat-Tek dishes (Mat-Tek Corporation). Glass substrates were coated with 20 µg/ml laminin (Invitrogen/Life Technologies).

#### HFF cell culture

Human foreskin Fibroblasts (HFF-1) were purchased from ATCC (ATCC® SCRC1041™) and cultured in medium consisted of high-glucose Dulbecco’s modified Eagle’s medium (all reagents from Life Technologies unless otherwise noted), 1 mM sodium pyruvate, 1× MEM-NEAA, 1% Pen/Strep, and 10% heat-inactivated FBS (Omega Scientific). Cells were passaged 1:5 with TrypLE every 4-5 days. For live cell TIRFM imaging, cells were plated on No. 1.5 glass Mat-Tek dishes (Mat-Tek Corporation) coated with 10 µg/ml human fibronectin (Corning).

#### MEF cell culture

Piezo1 heterozygous null mice (Ranade et al. 2014)were obtained from Jackson Laboratories (Stock No. 026948) and intercrossed to generate a mixture of wildtype, heterozygous, and knockout embryos. The time of the vaginal plug observation was considered E0.5 (embryonic day, 0.5). Dams were sacrificed E10.5, and mouse embryonic fibroblast (MEF) were derived from individual embryos as per Behringer et al. (Behringer et al. 2014). Tissue was mechanically dissociated, and cells were seeded onto plates coated with 0.1% Gelatin (Millipore ES-006-B), and passaged twice before experimentation. Growth medium consisted of DMEM (ThermoFisher, 11960-051), 10% FBS (Omega Scientific, FB-12), 1× GlutaMAX (ThermoFisher, 35050061), 1× Pen-Strep (ThermoFisher, 15140122), and 20 ng/ml PDGF (PeproTech, 100-00AB). Genotyping was performed through Transnetyx, and cells of the same genotype were pooled. Prior to TIRFM imaging, MEFs were dissociated with TrypLE Express (ThermoFisher, 12604013) and 5000 cells were plated onto 14 mm #1.5 glass Mat-Tek dishes coated with 10 µg/ml fibronectin (Fisher Scientific, CB-40008A). MEFs were imaged 2 days after seeding.

#### Generation of micropatterned square cells

Coverslips with square micropatterns were purchased from Cytoo (https://cytoo.com/, Catalog # CYTOOchips PADO-SQRS). CYTOOchips were coated with fibronectin per manufacturer’s instructions and cells were plated using a density of 1.5 × 10^4^ cells/ ml per manufacturer’s instructions in growth media. For TIRFM Ca^2+^ imaging live cells were imaged 2-5 hours after seeding. For immunofluorescence experiments, cells were fixed with 4% paraformaldehyde in phosphate-buffered saline supplemented with 5 mM MgCl_2_, 10 mM EGTA, 40 mg/ml sucrose, pH 7.5.

### Generation of Piezo1-knockout HFFs by CRISPR/Cas9

The Piezo1 gene was edited using the D10A nickase mutant of Cas9 (Cas9n) from S. pyogenes to limit off-target effects (Ran et al. 2013). The Zhang lab design tool:http://crispr.mit.edu/ was used to identify optimal and specific Guide A and Guide B sequences (Hsu et al. 2013). The guide sequences targeting Piezo1 exon 19 were cloned into plasmids with the sgRNA encoding backbone and had either the green fluorescence protein gene, 2A-EGFP (pSpCas9n(BB)-2A-GFP, PX461, Addgene Cat. #48140) or the puromycin resistance gene (pSpCas9n(BB)-2A-Puro (PX462) V2.0, PX462, Addgene Cat. #62987). PX461 and PX462 were a gift from Feng Zhang (Ran et al. 2013). Guide A sequence (GCGTCATCATCGTGTGTAAG) was subcloned into PX461 while Guide B sequence (GCTCAAG-GTTGTCAACCCCC) was subcloned into PX462.

Equal amounts of Guide A and Guide B plasmids (5 µg) were co-transfected into HFFs at passage 8 using NHDF Nucleofection® Kit (Neonatal cells protocol, Cat. # VAPD-1001 10) as per kit instructions using Nucleofector® Program U-020. Cells were treated with 5 µg/ml puromycin for 2 days following transfection (conditions in which all untransfected HFF cells die). Surviving cells were examined by fluorescence microscopy which revealed most cells to exhibit green fluorescence indicating that these cells contained both plasmids. Cells were plated to obtain single cells in 96-well plates (100 µl of 5 cells/ml per well) and expanded in 2% O_2_ and 5% CO_2_ incubator at 37º C. Genetic identification was performed by isolating gDNA from individual HFF clones using DNeasy Blood and Tissue kit (Qiagen) and amplifying the CRISPR/Cas9 targeted exon 19 region by PCR. The PCR products were sub-cloned into pGCBlue (Lucigen, pGCBlue Cloning and Amplification kit) or pMiniT (NEB PCR cloning kit, Cat. # E1202S) plasmids and sequenced. Sequence analysis of Clone 18-3 revealed out of frame indel modifications on both alleles in exon 19: 18 subclones had a 32 bp deletion with a 1 bp insertion (T), while 17 subclones had a 44 bp deletion.

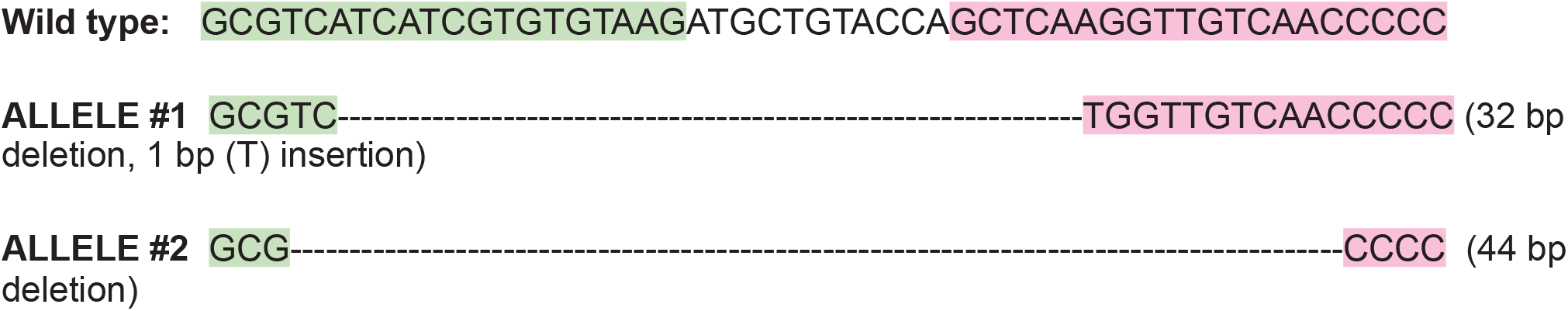

HFF clones were imaged in TIRFM assays as described above. As an appropriate control for experiments presented in Fig. 1B, a wildtype clone (9-7) isolated from the above procedure was used. We did not observe any differences in Ca^2+^ flickers in the parent HFF population and the 9-7 WT clone.

### Immunofluorescence Staining

Immunostaining was performed as previously described(Pathak et al. 2014) using the following antibodies: Rabbit anti-RFP (RFP Antibody Pre-adsorbed; Rockland, Cat# 600-401-379), 1:200 (0.95 μg/ ml) and mouse anti-paxillin (clone 5H11, Millipore Cat # 05-417), 1:1000, mouse anti-Integrin (IGTB1; clone 2B1, Fisher Scientific cat # MA10690), 1:100. Secondary antibodies used were Goat anti-rabbit Alexa Fluor 555 (Invitrogen Cat# A21428) and Donkey anti-mouse Alexa Fluor 488 (Invitrogen, Cat# A-21202), and Goat anti-Mouse Alexa Fluor 488 (Invitrogen, Cat# A11029) were used at 1:200 (0.01 mg/ml). Nuclei were stained by Hoechst 33342 (Life Technologies) at 4 μg/mL in PBS and actin filaments were stained with Phalloidin conjugated with TRITC (Sigma-Aldrich Catalog #P1951). Samples were mounted with Prolong Diamond anti-fade (Invitrogen cat # p36961).

### Western Blot

E12.5 Piezo1-tdTomato mNSPC spheres were lysed in RIPA buffer for 20 min on ice, then centrifuged for 10 min at 12,000 rpm at 4C. 30 μg protein was separated on Novex Wedgewell 6% Tris-Glycine gels (Life Technologies, cat #XP00060box) and subsequently transferred to PVDF Transfer Membrane, 0.45 µm (Invitrogen, Cat #88518). Piezo1-tdTomato was detected with anti-RFP antibody (1:1000, Life Technologies, Cat#: R10367) and 1:5000 Goat anti-Rabbit IgG (H+L) Secondary Antibody, HRP (Life Technologies, Cat# 31460) and detected with SuperSignal™ West Femto Maximum Sensitivity Substrate (Life Technologies, Cat# 34095) with Biorad ChemiDoc™ XRS+ system.

### Imaging

#### Imaging Piezo1 Ca^2+^ flickers

Piezo1 Ca^2+^ flickers were detected using Ca^2+^ imaging by TIRF microscopy. Cells were loaded by incubation with 1-2 μM Cal-520 AM (AAT Bioquest Inc.) in phenol red-free DMEM/F12 (Invitrogen) for 20-30 min at 37 °C, washed three times, and incubated at room temperature for 10–15 min to allow cleavage of the AM ester. Imaging was performed at room temperature in a bath solution comprising 148 mM NaCl, 3 mM KCl, 3 mM CaCl_2_, 2 mM MgCl_2_, 8 mM glucose, and 10 mM HEPES (pH adjusted to 7.3 with NaOH, Osmolarity adjusted to 313 mOsm/kg with sucrose). We refer to this solution as the standard imaging solution below.

Piezo1 Ca^2+^ flickers in Figs. 1, 2, 3, 6, S1, S2, S5, Movies M1, M2, M3 were imaged on a motorized Olympus IX83 microscope, equipped with an automated 4-line cellTIRF illuminator and a PLAPO 60× oil immersion objective with a numerical aperture of 1.45. Mat-tek dishes of shRNA-transfected cells loaded with Cal-520 AM were first scanned using an Olympus UPLSAPO 10× objective to identify cells expressing TurboRFP. Spatial coordinates of red fluorescent cells were marked using a programmable stage (Applied Scientific Instruments). Then the objective lens and illumination were switched for TIRF imaging, and previously identified red cells were imaged for Piezo1 Ca2+ flicker activity. Cells were illuminated with a 488 nm laser and images were acquired with a Hamamatsu Flash v4 scientific CMOS camera at 10 ms exposure and a frame rate of 9.54 frames/second.

Piezo1 Ca^2+^ flickers in hNSPCs in Fig. S3 were acquired at 200 Hz frame rate on a custom-built Spinning-Spot Shadowless TIRF microscope. Details of construction and comparison to traditional TIRF can be found in Ellefsen et al. 2015 (Ellefsen, Dynes, and Parker 2015).

An individual video is one microscope field of view, composed of one or more cells. Each experiment was performed multiple times, i.e. with cells prepared on different experiment days. On each experiment day we recorded from at least one but typically more than one dish of cells. Each video is unique, i.e. no cells were recorded multiple times (with the exception of Fig. 6D, where the same cells were imaged before and after Calyculin A treatment). Since cells have different cell spread areas, and cells in contact with each other can be hard to distinguish in live-cell images, we compute flicker frequency by unit area of the region covered by cells rather than per cell.

#### Imaging Piezo1 Ca^2+^ flickers and cellular traction forces in the same cell

Fabrication of Förster resonance energy transfer (FRET)-based molecular tension sensors (MTSs) to measure cellular traction forces was performed as previously described (Chang et al. 2016). The MTS is comprised of an elastic spring domain derived from spider silk, which is flanked by a covalently-bound FRET pair, Alexa 546 and Alexa 647. The N-terminus of the sensor possesses a HaloTag domain, while the C-terminal end presents the ninth and tenth type III domains of fibronectin.

Perfusion chambers (Grace Biolabs 622103) were attached to HaloLigand/PEG-functionalized coverslips. The MTS (at 0.03 mM for HFFs and 0.04 mM for hNSPCs) was added to the flow cell and incubated at room temperature for 30 min, washed with PBS twice, and passivated with 0.2% w/v Pluronic F-127 for 5 min. Flow cell channels were washed once with PBS before adding freshly dissociated cells in normal culture media and incubated at 37 °C with 5% CO_2_. Cells were typically allowed to spread for 1 h before imaging and not imaged for longer than 5 h after seeding. Cells were loaded with Cal-520 AM Ca^2+^ indicator as described above and imaged in DMEM/F12 medium containing 10% FBS and 3 mM CaCl_2_.

FRET-based traction force measurements and Piezo1 Ca2+ flicker measurements were performed with TIRFM on an inverted microscope (Nikon TiE) with an Apo TIRF 100× oil objective lens, NA 1.49 (Nikon). The FRET probe was excited with 532 nm (Crystalaser). Emission from Alexa 546 and Alexa 647 was separated using custom-built optics as described previously(Chang et al. 2016; Morimatsu et al. 2015). Donor and acceptor images were focused on the same camera chip. Data were acquired at 5 frames per second with an EMCCD camera (Andor iXon). Following imaging of the FRET force sensor, a motorized filter flip mount (Thor Labs) was used to switch emission filters for imaging Cal-520 Ca^2+^ indicator in the same cell. Cal-520 was excited using a 473 nm (Coherent Obis) laser and imaged at 15.29 ms exposure time.

#### Effect of pharmacological agents on Piezo1 Ca^2+^ flickers

ML-7 (Cayman Chemicals), Y-27632 (Sigma) and Calyculin-A (Cayman Chemicals) were dissolved in anhydrous dimethyl sulfoxide (DMSO) to make stock solutions of, 10 mM Y-27632, 50 mM ML-7 and 100 μM Calyculin A. Working concentrations used were 10 μM Y-27632, 50 μM ML-7, and 10 nM Calyculin A in standard imaging solution (see above for composition). For control measurements, comparable volumes of DMSO were added for each experiment. For experiments requiring 0 mM external Ca^2+^ the imaging solution used was 138 mM NaCl, 1 mM KCl, 5 mM MgCl_2_, 2 mM EGTA, 8 mM glucose, and 10 mM HEPES, pH 7.3, 313 mOsm/kg. For ML-7 treatment, cells were incubated in HFF Media containing 50 μM ML-7 for 30 minutes at 37 °C, then loaded and imaged with Cal-520 AM in the presence of 50 μM ML-7. For Calyculin A treatment, after control measurements in standard imaging solution, the bath solution was replaced with imaging solution containing 10 nM Calyculin A and cells were imaged after incubation for 1-5 minutes at room temperature.

#### Imaging Piezo1 diffusion with TIRFM

For Piezo1 diffusion studies in Fig.5, images were acquired on a Nikon N-STORM system built around a Nikon Eclipse Ti microscope. The imaging objective used was a Nikon 100× APO TIRF oil immersion objective (NA 1.49). Images were acquired on an Andor iXon3 electron-multiplying charge-coupled device (EMCCD) camera with an 100 ms exposure time and 160 nm/px in TIRF mode. Cells were continuously illuminated with a 561 nm laser.

#### Confocal imaging

Confocal imaging was performed on a Zeiss Confocal Spinning Disc Confocal Microscope (Zeiss) using a 63X objective with a numerical aperture of 1.40. Image stacks were acquired with 405nm, 488nm, and 561nm lasers, in intervals of 0.3 µm thickness using the AxioVision Rel 4.8 software.

### Image analysis

#### Automated detection of Piezo1 Ca^2+^ flickers

Piezo1-mediated Ca^2+^ flickers were detected using an improved version of our published algorithm for automated detection of Ca^2+^ signals(Ellefsen et al. 2014). The new algorithm, which runs as a plug-in under the open-source image processing and analysis package Flika (https://github.com/flika-org/flika), uses a clustering algorithm(Rodriguez and Laio 2014) to group super-threshold pixels into calcium events, improving both signal detection and segregation of signals which overlap temporally or spatially.

An F/F_0_ movie is generated from the original recording by subtracting the camera black level and dividing each pixel at every frame by its average value across the first ~100 frames. To remove low temporal frequency signal drift, the F/F_0_ movie is temporally filtered with a high pass Butterworth filter. To standardize variance across pixels, the value of each pixel is divided by the standard deviation of the values at baseline. The noise in this ‘normalized’ movie is normally distributed with a mean of 0 and standard deviation of 1.

A threshold is applied to a spatially-filtered version of the ‘normalized’ movie to generate a binary movie. Each super-threshold pixel in this binary movie is putatively considered part of a flicker. In order to group these pixels together, we modified the clustering algorithm published by Rodriguez and Laio(Rodriguez and Laio 2014). Briefly, a density is assigned to every super-threshold pixel by counting the number of pixels in an user-defined ellipsoid centered around the pixel. Then, for every pixel, the distance to the nearest pixel with a higher density is determined. Pixels that represent the center of clusters will have both a high density and a high distance to a pixel with higher density. The user manually selects pixels exceeding a density and distance threshold as cluster centers. The algorithm then assigns every other pixel to a cluster center pixel recursively, by finding the cluster of the nearest pixel of higher density. Once all pixels have been clustered, clusters below a user-defined size are removed.

After flickers have been identified by the clustering algorithm, the subpixel centroid of the signal is found by averaging each pixel in the ‘normalized’ movie over the flicker duration, and fitting a 2D Gaussian function to this average image. The peak amplitude, temporal dynamics, and frequency of signals at specific sites can be quantified, and the resulting data can be exported as Excel or csv files.

This algorithm is implemented in the puff_detect plugin for the image analysis software Flika, downloadable at https://github.com/kyleellefsen/detect_puffs. Both the puff_detect plugin and flika are open source software written in the Python programming language. Instructions for installation and use of the algorithm can be found at http://flika-org.github.io/.

#### Generation of cellular force maps

Analysis of FRET signals from the MTS was performed following the methodology from Morimatsu & Mekhdjian et al., Nano Letters 2015. Briefly, FRET index maps were generated by dividing the acceptor intensity A (background subtracted) by the sum of the acceptor and donor (D) intensities (also background subtracted): FRET_i_ = A / (A + D). FRET index maps can be converted to FRET efficiency maps to extract quantitative values for force from the FRET efficiency to force calibration curve. FRET index is converted to FRET efficiency using the following equation:

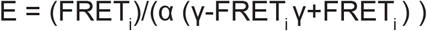

Where E is the FRET efficiency, FRET_i_ is the FRET index, α is the fraction of donor-labeled sensors that have an acceptor, and γ is a factor that accounts for differences in donor and acceptor quantum yield. Both α and γ are experimentally determined (Morimatsu et al., Nano Letters 2015). The FRET efficiency is converted to force using a phenomenological fit (Chang et al., ACS Nano 2016) to the FRET-force response of the (GPGGA)_8_ linker (Grashoff et al., Nature 2010).

#### Calculation of distance from Piezo1 Ca2+ flicker localization to nearest force-producing region

Force-generating regions were determined by blurring the force maps with a Gaussian filter. Regions in which the pixel intensity was below 75% of maximum intensity were considered force generating. Distances from each flicker centroid to the nearest force generating region were measured. To calculate the average distance to the nearest force generating region in each cell, the outline of each cell was manually traced, 1000 points were randomly selected inside this outline, and the distance to the nearest force generating region was measured.

#### Piezo1 particle tracking

TIRFM image stacks were processed in order to determine the location of Piezo1-tdTomato puncta in each frame. Each frame was spatially bandpass filtered by taking the difference of Gaussians, an image processing algorithm that enhances a band of spatial frequencies--in this case, around the size of the particles. The spatially filtered movie was then thresholded using a manually determined threshold, yielding a binary movie. Spatially contiguous pixels above threshold were grouped together and considered a single particle. The centroid for each particle was determined by fitting a 2D Gaussian function to each particle, yielding a centroid with subpixel precision. The initial x, y values for the fit were set to be the center of mass of the binary pixels in the particle. Any localizations within consecutive frames that were within three pixels of each other were assumed to arise from the same particle. These localizations were linked over time to generate particle tracks.

### Data analysis and statistical testing

Data generated or analyzed during this study are included in this article (and its supplementary information files) along with detailed methods, descriptions and sample movie files where appropriate. OriginPro 2018 (OriginLab Corporation) was used for statistical analysis and generating plots. P values and statistical tests used are indicated in figure legends. A two-sample t-test was used where data were modeled by a normal distribution and the non-parametric Kolmogorov-Smirnov test was used in the case of non-normal distributions.

---

**Figure S1.**
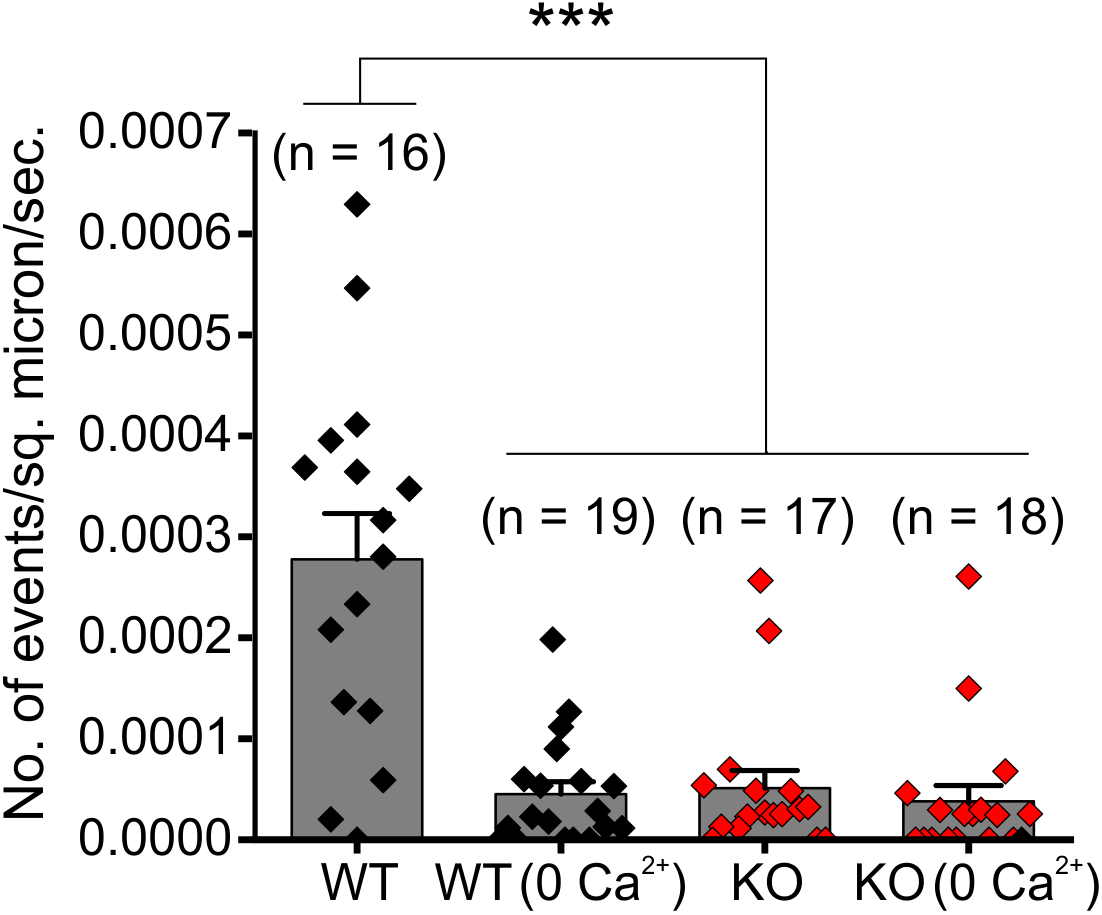
Residual Ca^2+^ flickers in Piezo1-knockout HFFs are mediated by intracellular Ca^2+^ release. Related to Fig. 1. Ca^2+^ flickers in WT HFF cells are reduced in Ca^2+^-free extracellular solution (denoted as “0 Ca^2+^”). Piezo1knockout HFF cells in standard 3 mM extracellular Ca^2+^ exhibit Ca^2+^ flickers to a similar extent as WT (0 Ca^2+^) cells, and these residual flickers are not eliminated in Ca^2+^-free extracellular solution. Taken together, these observations suggest that residual Ca^2+^ flickers in Piezo1-knockout HFFs occur due to intracellular Ca^2+^ release rather than plasma membrane ion channels. *** p < 0.001 by Kolmogorov-Smirnov test.

**Figure S2.**
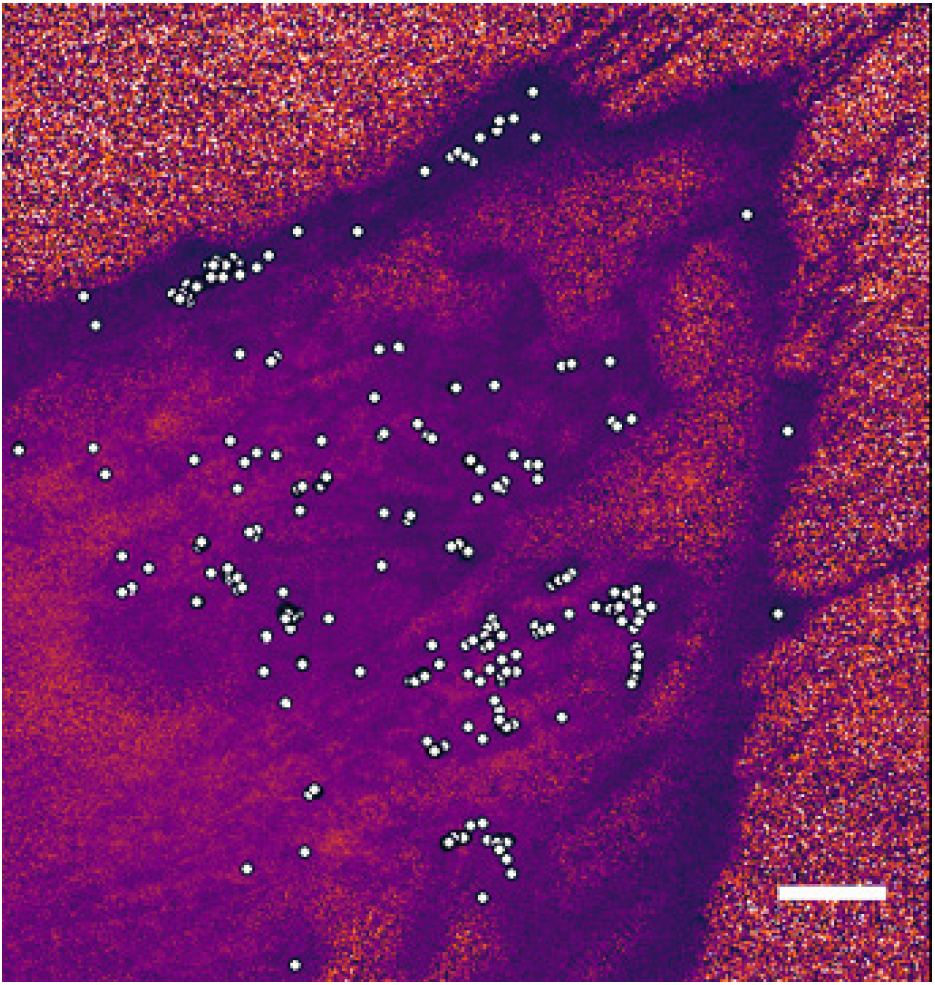
Piezo1 flicker localizations extracted from Movie S1. Related to Fig 2. Piezo1 Ca^2+^ flickers from HFF Movie M1 were analyzed as described in Fig. 2 and Methods. Scale bar, 10 µm.

**Fig. S3.**
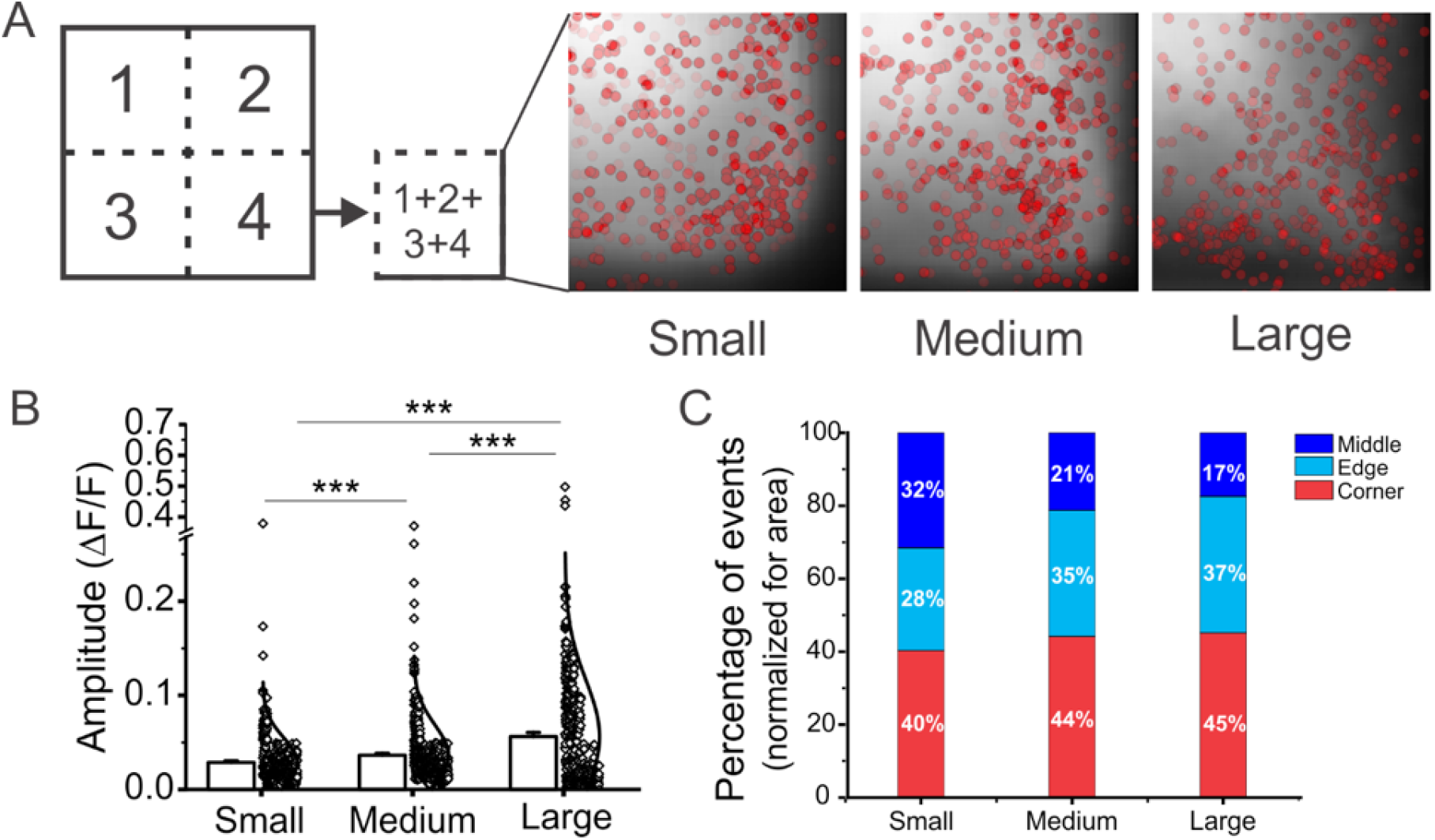
Enrichment of Piezo1 Ca^2+^ flickers in Corners and Edges in square hNSPCs. Related to Fig. 3. **A.** Each panel shows a quadrant of an Cal520-loaded HFF, with an overlay of Piezo1 flicker localizations (red dots); since the square is a symmetric shape, flicker localizations from all four quadrants of cells are displayed on a single quadrant. Total numbers of events represented in panels C, D, E are: Small, 453 events from 7 cells; Medium, 476 events from 7 cells; Large, 426 events from 4 cells. The panels are shown scaled to the same size for ease of comparison. **B.** Bar and Data plot of flicker amplitudes (ΔF/F) for events from cells adhering to Small, Medium, and Large islands. Each dot represents an individual Piezo1 flicker event and bars denote the mean of all events for the specified cell size. Piezo1 Ca^2+^ flickers show greater amplitudes in cells seeded on large islands. *** denotes p < 0.001 by Kolmogorov-Smirnov test. **C.** Distribution of events in Corners, Edges and Middle regions for cells seeded on Small, Medium and Large squares. Difference of distribution patterns in hNPSCs and HFFs (see Fig. 3) suggests that the Piezo1 channel is tuned to different operating ranges in the two cell types. Chi-square test results: for Small cells χ^2^(2, N = 453) = 11.98, p < 0.01; for Medium cells χ^2^(2, N = 476) = 27.979, p < 0.0001; Large cells, χ^2^(2, N = 426) = 38.113, p < .0001.

**Figure S4.**
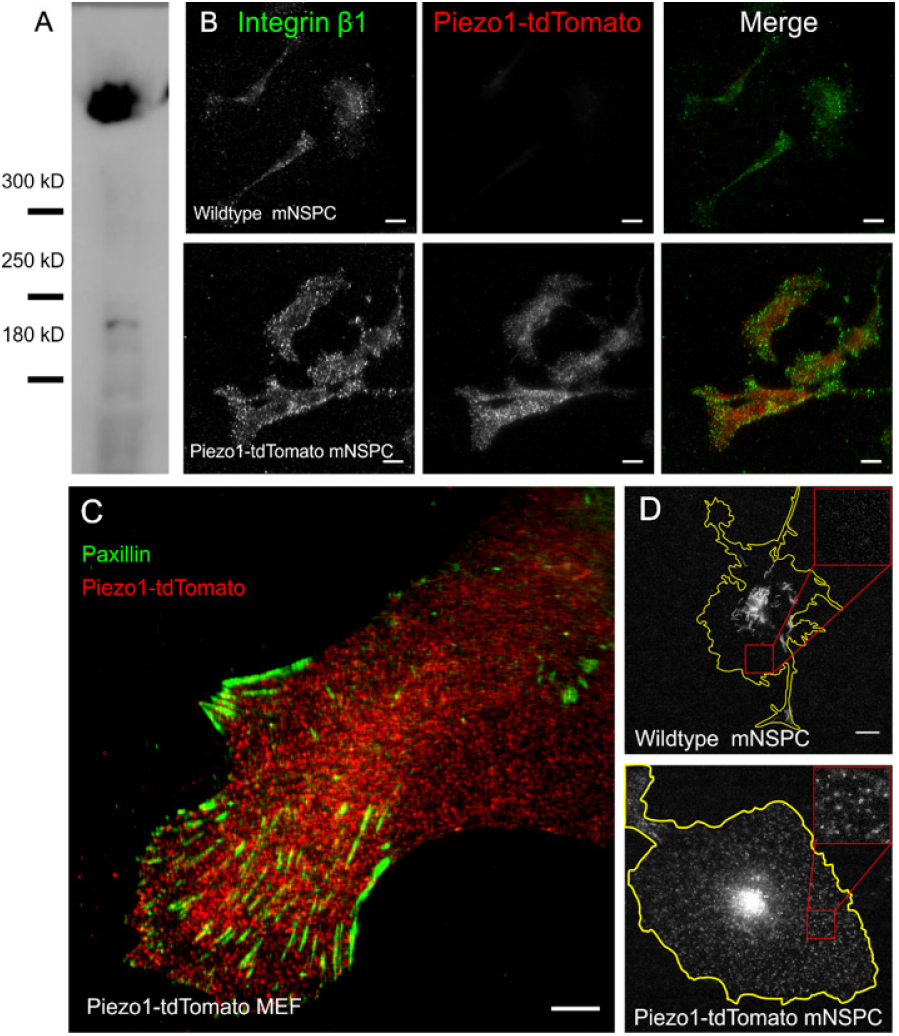
Piezo1 localization is not restricted to integrin-rich focal adhesions. Related to Fig. 5. **A**, Western blot of lysate from Piezo1-tdTomato mNSPCs probed by an anti-RFP antibody shows a band of the molecular weight expected for a Piezo1 and tdTomato fusion protein (340 kD). **B**, Representative TIRFM images of mNSPCs immuno-labeled with antibodies against Integrin β1 (green) and tdTomato (red). The top row shows images of mNSPCs harvested from wildtype mice, and the bottom row shows images of mNSCPs harvested from Piezo1-tdTomato reporter mice. Scale bar = 10 µm. **C.** Representative TIRFM images of MEFs immunolabeled with antibodies against focal adhesion protein paxillin (green) and tdTomato (red). Scale bar = 10 µm. **D.** Representative TIRFM images of tdTomato fluorescence of live mNSPCs from wildtype mice (top) and from Piezo1-tdTomato reporter mice. Inset shows magnification of a region of interest. While both cell types show autofluorescence (most prominent in the nuclear region), only Piezo1-tdTomato cells show small distributed puncta of tdTomato fluorescence. Scale bar = 10 µm.

**Figure S5.**
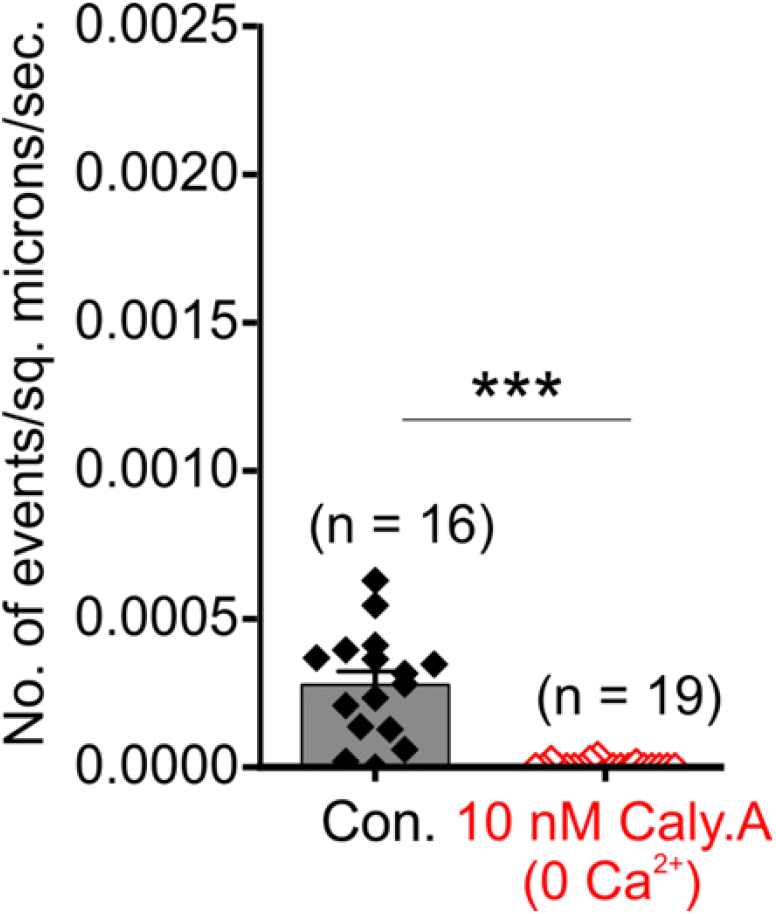
Increase in Ca^2+^ flickers with Calyculin A requires external Ca^2+^. Related to Fig 6. Flicker frequency in Control imaging solution and 1-5 minutes after replacing the control bath solution with 10 nM Calyculin A in Ca^2+^-free imaging solution (i.e. standard imaging solution containing 2 mM EGTA and no CaCl_2_). Bars denote Mean ± sem and each point represents flicker frequency in an individual video. Data are from three experiments. *** denotes p < 0.001 by Kolmogorov-Smirnov test.

## Movies

**Movie M1. Piezo1 Ca^2+^ flickers are reduced in Piezo1-knockout HFFs.** The movie shows an F/F_0_ ratio movie of Ca^2+^ flickers from WT and Piezo1-KO HFFs.

**Movie M2. Piezo1 Ca^2+^ flickers are reduced in Piezo1-knockout MEFs.** The movie shows an F/F_0_ ratio movie of Ca^2+^ flickers from WT and Piezo1-KO MEFs.

**Movie M3. Piezo1 Ca^2+^ flickers imaged from HFFs.** The movie shows the F/F_0_ ratio movie from WT HFFs that was used for localization of Piezo1 Ca^2+^ flickers in Fig. S2.

**Movie M4. Mobility of Piezo1-tdTomato puncta in mNSPCs imaged using TIRFM.** Puncta visible outside the yellow cell line are from neighboring cells.

